# Evidence of independent acquisition and adaption of ultra-small bacteria to human hosts across the highly diverse yet reduced genomes of the phylum Saccharibacteria

**DOI:** 10.1101/258137

**Authors:** Jeffrey S. McLean, Batbileg Bor, Thao T. To, Quanhui Liu, Kristopher A. Kerns, Lindsey Solden, Kelly Wrighton, Xuesong He, Wenyuan Shi

## Abstract

Recently, we discovered that a member of the Saccharibacteria/TM7 phylum (strain TM7x) isolated from the human oral cavity, has an ultra-small cell size (200-300nm), a highly reduced genome (705 Kbp) with limited *de novo* biosynthetic capabilities, and a very novel lifestyle as an obligate epibiont on the surface of another bacterium ^1^. There has been considerable interest in uncultivated phyla, particularly those that are now classified as the proposed candidate phyla radiation (CPR) reported to include 35 or more phyla and are estimated to make up nearly 15% of the domain Bacteria. Most members of the larger CPR group share genomic properties with Saccharibacteria including reduced genomes (<1Mbp) and lack of biosynthetic capabilities, yet to date, strain TM7x represents the only member of the CPR that has been cultivated and is one of only three CPR routinely detected in the human body. Through small subunit ribosomal RNA (SSU rRNA) gene surveys, members of the Saccharibacteria phylum are reported in many environments as well as within a diversity of host species and have been shown to increase dramatically in human oral and gut diseases. With a single copy of the 16S rRNA gene resolved on a few limited genomes, their absolute abundance is most often underestimated and their potential role in disease pathogenesis is therefore underappreciated. Despite being an obligate parasite dependent on other bacteria, six groups (G1-G6) are recognized using SSU rRNA gene phylogeny in the oral cavity alone. At present, only genomes from the G1 group, which includes related and remarkably syntenic environmental and human oral associated representatives^1^, have been uncovered to date. In this study we systematically captured the spectrum of known diversity in this phylum by reconstructing completely novel Class level genomes belonging to groups G3, G6 and G5 through cultivation enrichment and/or metagenomic binning from humans and mammalian rumen. Additional genomes for representatives of G1 were also obtained from modern oral plaque and ancient dental calculus. Comparative analysis revealed remarkable divergence in the host-associated members across this phylum. Within the human oral cavity alone, variation in as much as 70% of the genes from nearest oral clade (AAI 50%) as well as wide GC content variation is evident in these newly captured divergent members (G3, G5 and G6) with no environmental relatives. Comparative analyses suggest independent episodes of transmission of these TM7 groups into humans and convergent evolution of several key functions during adaptation within hosts. In addition, we provide evidence from *in vivo* collected samples that each of these major groups are ultra-small in size and are found attached to larger cells.

## Introduction

The phylum TM7, and more recently proposed name, Saccharibacteria, has been an enigmatic phylum of bacteria due to the fact that it had remained uncultivated for more than two decades since its first SSU ribosomal sequence (^2^ and recognition that is a candidate phyla ^3^. Over the years it appeared cosmopolitan in its distribution, inhabiting many environmental locations and within insect and mammal hosts. We recently discovered ^1^ through directed cultivation from human oral samples, that a member of this phylum (formal designation *Nanosynbacter Lyticus* Type Strain TM7x HOT 952) has an extremely reduced genome, ultrasmall cell size and lives as an obligate epibiont on the surface of a partner oral species, a subspecies of *Actinomyces odontolyticus ^4^*. This member of TM7 also exhibits what has been defined as a parasitic phase where it disrupts the bacterial host cell and causes death but remains viable itself to re-infect when a fresh host is available resulting in a stable co-culture early on. This discovery and co-culture marked the first concrete evidence of how these ultra-small organisms survive and is beginning to reveal a deeper understanding the host bacterium dependence and dynamics ^5^.

The phylum members from the G1 group to date all have extremely reduced genome size overall ^6^ ^7^ and the oral representatives approach the smallest known genomes of host associated microorganisms including Mycoplasma and obligate intracellular bacteria such as those within insect hosts (*Buchnera*, *Wigglesworthia* and *Blochmannia*). TM7 was only recently revealed to be part of the CPR when a larger study using filtered DNA from groundwater revealed many new candidate phyla formed a distinct group using phylogenetic markers (Figure 1) ^8,9^. The CPR is estimated to comprise 15% of the Domain Bacteria with upwards of 35 phyla by some estimates (Figure 1A and B) ^8^. To date, TM7x represents the only member of this entire CPR that has been cultivated and fully sequenced with a stable co-culture system on its host bacteria and therefore represents the type strain for this phylum. Currently, only three CPR are currently routinely detected in the human body (GN02, SR-1, and TM7). SR-1 are widely distributed in the environment and hosts similar to TM7 with one near complete genome from the oral cavity ^10^, yet has no cultivated representatives to date. Less is known about phylum GN02 and no human associated genomes from this group are yet available.

**Figure 1.**
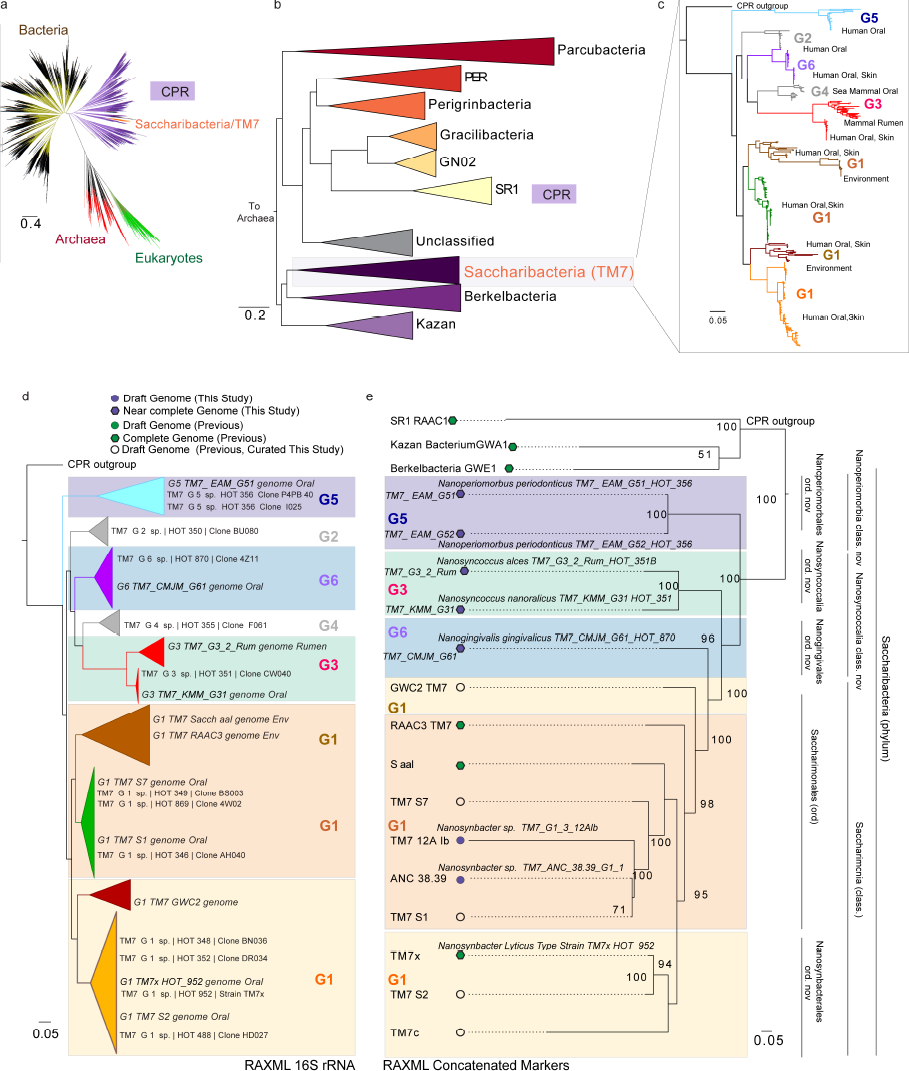
Phylogeny of the Saccharibacteria (TM7). (**a**) Current view of the tree of life highlighting the Saccharibacteria and Candidate Phyla Radiation (tree inferred from concatenated ribosomal gene dataset provided by Hug et al. 2016) (**b**) Phylum level maximum likelihood concatenated ribosomal gene tree of the CPR indicating Saccharibacteria share a common ancestor with Berkelbacteria and Kazan groups. (**c**) Fast tree 16S rRNA gene phylogeny of the major Saccharibacteria groups derived from public ribosomal databases and available genomes. The complete tree is available in circular format with full bootstrap values as Extended Data Figure 1 and in newick format in Supplementary Dataset 1. (**d and e**) Phylogenetic relationships within the Saccharibacteria phylum using maximum likelihood 16S rRNA and concatenated ribosomal marker gene inference. (d) Maximum likelihood 16S rRNA gene phylogeny and shown in c, sequences from Human Oral Microbiome Database and extracted from genomic assemblies shown. (**e**) concatenated ribosomal RAXML gene tree for new and previously published complete and draft genomes including first representative sequences from the G5 (*Candidatus* Nanoperiomorbus *periodonticus, class. nov.*), G6 (*Ca.* Nanogingivalis *gingivalicus ord. nov.*) G3 (Ca. Nanosyncoccus *nanoralicus, Ca. Nanosyncoccus alces. ord nov*) groups. Genome properties and proposed names for groups within the novel class, and orders proposed explained in Supplementary Table 1 and 2. CPR phyla SR1, Kazan, and Berkelbacteria used as outgroups.

Many SSU ribosomal marker gene studies have revealed the broader TM7 diversity in environmental ^3^ and human oral ^11^ and skin samples. Camanocha et al. ^12^ provided a comprehensive analysis of the phylogeny of the CPR groups in the oral cavity using phyla selective PCR primer pairs. From the cumulative curation and phylogenetic analysis of the 16S rRNA gene sequences available at this time six distinct clade level TM7 sequences are proposed. The original designations of these sequences include a Human Oral Taxa (HOT) ID designations G1 through G6 ^12^. Overall, TM7 members have been detected in various human body sites ^13-17^ and its association with inflammatory mucosal diseases has been implicated ^13,17,18^. The phylum is particularly prevalent in the oral cavity in and has been detected in Neanderthal ^19^ and ancient dental calculus of hunter-gatherers ^20^. Although commonly at low abundance, generally around 1% of the oral microbial population based on culture independent molecular analysis ^13,21^, an increase in abundance (as high as 21% relative abundance in some cases) of TM7 members was detected in patients with various types of periodontitis ^22,23^. Furthermore, specifically G5 and certain G1 (clone AY134895) are more prevalent in diseased samples and some of these can even be detected on or within the host crevicular epithelial cells ^24^. Based on these findings, the association of TM7 with diseases such periodontitis is evident. With a single copy of the 16S rRNA gene resolved on a few limited genomes, which is also consistent for most CPR^8^, their absolute abundance is therefore most often underestimated and their potential role in disease pathogenesis could therefore be more important than previously recognized. Currently, 16S rRNA gene surveys and metagenomics studies have not yet resolved the members in these samples to the taxonomic level of the groups (HOT) due to the relatively recent establishment of phylogenetic diversity with full length 16S rRNA sequences in databases in the former case as well as a complete lack of genomes outside of the G1 group in the latter approach. Recognizing there is an immediate need for increased resolution for identification, understanding niche specialization linked to specific disease sites as well as the encoded virulence potential of each of the groups, we systematically sought to capture reference genomes from all the host associated groups and also develop a naming convention of these new assemblies to link Saccharibacteria genomes to Human Oral Taxon designation from 16S rRNA genes. Here we report on novel genomes covering the spectrum of known diversity enabling insights into the adaptations that have occurred from environment to living within hosts, the evolution, ecology and overall genomic variation as well as the sub micron physical cell size variation across this very unique phylum.

## Results

### Diversity of the Phylum Saccharibacteria beyond the G1 group

Current knowledge of the diversity in the Saccharibacteria/TM7 phylum has been limited to 16S rRNA gene sequences and genomes from the G1 group. Using available 16S sequences from HOMD and newly extracted 16S sequences from genome assemblies, (Figure 1c, d) the inferred SSU tree reveals the G1-G6 clades are monophyletic with branching subclades (Extended Data Figure 1 (Full 16S tree), Supplemental Dataset as newick tree file). Major sources of the sequences are highlighted in the condensed tree showing the mixed environmental and host associated sources in the G1 group observed previously, and intriguingly mammalian rumen and human oral sources forming distinct clades in the G3 group. We recovered genome bins for these uncultivated, typically low abundance groups G3, G6 and G5 as well as additional G1 bins from a combination of approaches (mainly binning from metagenomic data) and sources (see Supplemental Tables 1-3 for summary of genome bin sources, curation, and properties). We then investigated their phylogenomic relatedness using a concatenation of 42 ribosomal marker genes with other CPR included as outgroups (Figure 1) and proposed operational taxonomic classifications (Supplemental Table 2) based on these inferred maximum likelihood trees and through the comparative genomic analysis described throughout this study including 16S rRNA and genome based average amino acid identities. The 16S rRNA gene tree (Figure 1d) was highly congruent with the marker gene tree for the genomes (Figure 1e). We derived the most distantly related G5 group genome bins as well as a G1 member from metagenome samples from patients ^25^ with severe periodontitis (*Nanoperiomorbia* Class nov.). Sequencing and genome binning of lab enrichment cultures from oral derived samples, similar to the approach for isolating TM7x, resulted in the G3 oral genome bin. Mining a metagenomic sample derived from moose rumen revealed a genome for the related G3 rumen that formed a distinct clade using ssu rRNA next to the oral G3 (*Nanosyncoccalia* Class Nov.), confirming the presence of this group in mammalian rumens. Human Microbiome Project (HMP) datasets (1149 assemblies 15 body sites, ~3.5 Tb) were scanned for 16S rRNA gene hits to CPR those samples with relatively long contigs were then re-assembled and binned which yielded a G6 genome bin from a keratinized gingiva sample (*Nanogingivales* Ord. Nov). An ancient G1 group representative genome bin was also derived from dental calculus using whole DNA shotgun methods from adult human skeletons (B61 skeleton) at the medieval monastic site of Dalheim, Germany (c. 950–1200 CE) with evidence of mild to severe periodontal disease ^26^. Based on the phylum name Saccharibacteria, we propose the Class Saccharimona which includes published G1 genomes from environmental sources including deep subsurface groundwater and sludge bioreactor as well as oral genomes from single cell approaches and cultivated strain TM7x, forming two possible Orders, *Saccharimonales* containing both oral and environmental representatives as well as *Nanosynbacterales* (Ord Nov.) which is monophyletic and populated with oral genomes at this time. However the environmental genomes might possibly not be a coherent group (GWC2 separated and S. aalborgensis grouped with RAAC3), also supported by the concatenated gene tree. We noted that G2 and G4 sequences were not found in the HMP metagenomes on contigs of significant length and therefore did not result in any genome information. This is consistent that the sequences from these groups found less often in databases indicating they may be extremely low abundance but G4 may be enriched for in sea lion and dolphin oral cavity where several hits were reported ^27,28^. Overall we have uncovered substantial sampling of the known phylum diversity with representative genomic datasets that span environmental and host-associated members allowing further exploration of this unique CPR group.

### Human biogeography and ecology from Neanderthal to modern humans

Saccharibacteria, SR1 and GN02 are unique among the CPR in that they have transitioned from the environment to live in mammalian hosts. With this new genomic and phylogenomic information for the Saccharibacteria and oral SR1 genome, we sought to re-investigate the human biogeography and ecology of these Phyla from Neanderthal to modern humans by mapping shotgun reads from available datasets as well as SSU RNA from assemblies and amplicon datasets. Neanderthal dental calculus, estimated at 48 thousand years (kyr), indicates G5 and many G1 can be readily detected with little to no coverage of G3, G6 or SR-1 (Figure 2a, Supplementary Table S4). Coverage statistics of TM7x, used as an oral reference genome, was reported in the original study. Increased genome coverage is evident in medieval dental calculus (~1200CE) including G3 and SR1. Metagenomic data confirms different groups are found in varying abundances across body sites within just these representative individual samples. Analyses of the 16S rRNA sequences amplified from the HMP subjects have also previously revealed the distribution of TM7 at the phylum level resolution given the known sequences in databases at the time. Estimated abundances across 10 body sites of the human digestive tract with proportions of hits to the phylum ranging from 0.13% to 5.7% in the oral cavity sites and .008% in stool ^29^. Oligotyping analyses of HMP datasets has also reported distribution of TM7 sequences at single nucleotide resolution across the oral cavity and stool ^30^. Re-analyses of this dataset provides additional information in terms of phylum proportions using the best hit to TM7 groups across 10 oral cavity sampling sites (Extended Data FIGURE 2). We then chose to confirm each CPR groups prevalence in significant abundance within the HMP assemblies as amplicon sequencing depth is higher but provides lower phylogenetic resolution due to the typical use of short reads. TM7 and CPR groups (FIGURE 2b, Supplementary Table S4) were identified across body sites within contigs (>300bp) containing the ssu rRNA gene. We show that the TM7, GN02, SR-1 distributions are dominant in the oral cavity with few hits in other body sites. Our G5 group genome was obtained from subgingival pocket of a periodontitis patient and had previously been associated with this disease due to their drastic abundance increase. We confirmed that group is also predominately found enriched in subgingival pockets within healthy subjects. High coverage of G1 groups in Neanderthal and Medieval samples is consistent with the supragingival plaque from which mineralized calculus is formed.

**Figure 2.**
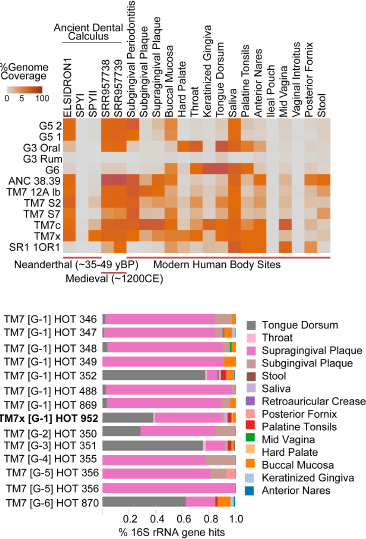
Ancient to Modern Biogeography and Niche Specialization. (**a**) Percentage of genome coverage from mapped metagenomics read sets to newly available Saccharibacteria groups from Ancient Neanderthal dental calculus (48,000 years), through to modern body sites in health and disease. Neanderthal calculus from the best-preserved Neanderthal, El Sidrón 1, which suffered from a dental abscess is included as well as well as SPYNEWWL8 which had extensive DNA damage (Supplemental Table 4) (**b**) Proportional distributions of the distinct Saccharibacteria groups from 16S rRNA gene hits within body site specific assembled contigs from the Human Microbiome Project (>300bp hit cutoff, n=1100 assemblies) highlighting the niche specialization across the groups (Extended Data Figure 2 with supporting oligotyping data additional SR-1 and GN02 group distributions).

Outside of the human digestive tract, there is evidence for TM7 phylum in the skin (retroauricular crease), mid vagina and stool but at low prevalence, which may reflect the number of assemblies, depth of sequencing but most likely due to the low relative abundance of these groups in other body sites. TM7 G5 was notably high in the posterior fornix of the vagina as were hits to GN02 with one SR-1 (>300bp) in the mid-vagina bodysite (Extended Data Figure 2) confirmed with metagenome mapping showing their presence but lower abundance in the non-oral sites. Overall there are variations in the types and proportions of Saccharibacteria, SR1 and GN02 across the human body sites depending on the approach but 16S amplicon studies generally agrees with the results we obtained screening the HMP assemblies and metagenome mapping that all support the specific as well as overlapping ecological niches that each of the groups can occupy.

### Ultrasmall cell size are a common feature of the Phylum Saccharibacteria and SR1 in the human oral cavity

Fresh human oral saliva and tongue-surface bacterial samples were collected and screened for the presence of TM7-clades in 12 individuals using PCR. Two individuals showing positive identification for TM7-G3, G5, G6 and SR1 were studied further. The samples were either stained with clade specific fluorescence in situ hybridization (FISH) DNA probes that specifically designed to target the 16S rRNA sequence of the bacteria, or filtered through 0.45 um membrane to select for the ultra-small bacteria. Previous studies showed that TM7 specific FISH probes could be non-specific and stain other bacteria. To insure specific staining, we designed our probe carefully by looking at our novel genome information, as well as performed FISH analysis on TM7x (G1) strain ^1^ using these TM7-clade specific probes, and showed that all the probes except G1-specific probes do not stain TM7x cells (Methods). Results of FISH on these *in vivo* derived samples clearly showed that TM7 (G1, G3, G5 and G6) and SR1 are small bacteria that decorate larger-host bacteria, similar to our isolated TM7x (Figure 3). In addition, filtered samples were positive for the presence of G3, G5, G6 and SR1 groups (Figure 3), supporting the imaging data that these bacteria generally have ultra-small cell sizes. These results are entirely consistent with their shared phenotypes having reduced genomes and limited *de novo* biosynthetic capacity. Initial screening for G2 and G4 using clade specific probes revealed that no positives in our sample collections further supporting that these genomes are more rare in the human oral cavity and/or are mainly restricted to other hosts and have been misidentified in oral samples. Overall the FISH imaging and new genomic information from G3 G5 and G6 are also highly supportive of an obligate symbiotic lifestyle like strain TM7x including the reduced genomes, the missing capacity of *de novo* biosynthesis of amino acids and vitamins.

**Figure 3.**
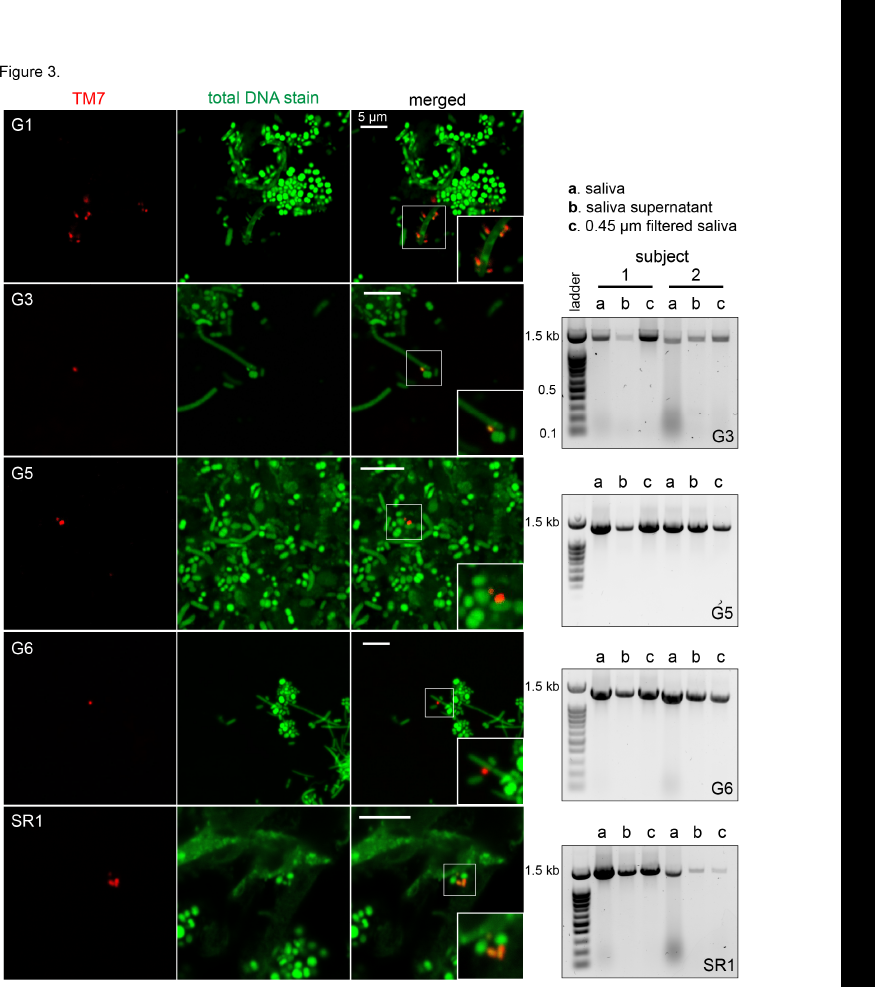
Visualization of different TM7 groups and SR1 in saliva samples. **(a)** Human saliva and ton Figure3_Batmarch20_v2-01gue samples were stained with *Fluorescence In Situ Hybridization* (FISH) probes that specifically targeting G1, G3, G5 and G6 clades of TM7 phylum, as well as SR1 phylum. Column one is TM7 staining with clade specific probes (red), column two is total DNA stain with Syto9 dye (green), and column three is merged images (magnified images indicate ultra-small bacteria). All scale bars are 5 µm. (b) PCR amplification of G3, G5, G6 and SR1 16S rRNA with clade specific primers, using template DNA isolated from original saliva samples, saliva supernatants and filtrate passing through 0.45 µm filter (see methods) confirming ultra-small cell sizes.

### Reduced genomes from environment to host with specific convergent host adaptations across host associated groups

Genome wide pairwise average amino acid identity (AAI) analysis between orthologous genes and pangenomic comparisons resulted in a 2906 ortholog gene set (11,615 total genes, 15 TM7 genomes) along with close CPR (SR1, KAZAN and Berkelbacteria) as outgroups, highlight the surprisingly large genetic variation across this phylum (Figure 4, Extended Data Figure 3). These analyses provided further support of the phylogenetic relationships between the groups observed with SSU and concatenated gene approaches both with a set of single copy ribosomal markers (Figure 1) as well as using a concatenated set of 121 core genes and overall gene presence/absence (Extended Data Figure 3). Comparing representative genomes of each group with the environmental G1 genomes, a set of 201 core genes are shared (Figure 4, Supplementary Table S6), 201 between oral and environmental (777 unique oral genes) and 208 are between just the HA oral groups. The Saccharibacteria pangenome appears to be open at this stage of genomic discovery. G5 *Nanoperiomorbia* (Class nov.) has less than 53% AAI with any other genome, shares only 45% of its genes with other groups.

**Figure 4.**
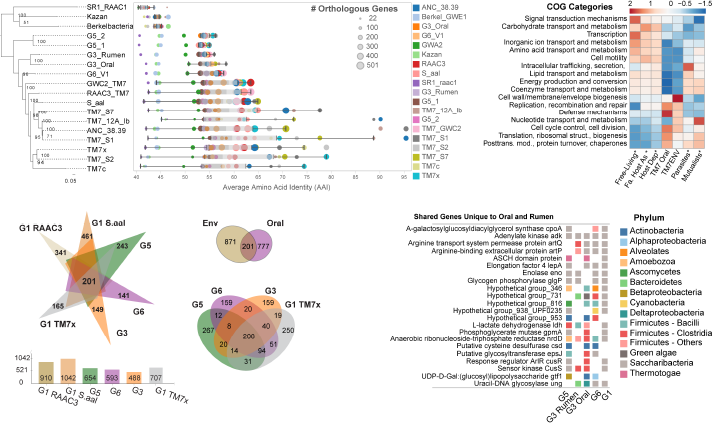
Divergent human associated Saccharibacteria share reduced genomes and acquired genes in a convergent manner for host adaptation. (**a**) Relationships between phylogeny with average amino acid identities and number of shared orthologous genes between Saccharibacteria and other divergent phyla highlight the low number of orthologous genes and percent homology between groups in these reduced genomes (percent identity cutoff >=20%, Full table available in Supplemental Data Figure 1) (**b**) COG distribution percentage across genomes for functionally annotated genes within free-living, facultative host-associated and host dependent as well as obligate intracellular (mutualistic and parasitic bacteria) (derived from Merhej et al. (20)) compared to environmental and oral Saccharibacteria groups. Enrichment of genes with functions in defense mechanisms, translation, replication/recombination and lack of those involved with amino acids, lipid and energy production and conservation supporting the reduced functional capacity and host dependent lifestyle (Supplemental Table 5; Extended Data Figure 5). Rows are centered and unit variance scaled. (**c**) Pangenome analysis of environmental and host associated groups reveals the large number of unique genes in each new genome with a core of 201 genes. Oral genome comparisons using the available genomes from each group share 208 core genes with unique genes ranging form 159 to 267. (Supplemental Table 6 and 7) (**d**) A total of 21 unique genes are shared with 4 or more of the oral genomes and are not found in environmental genomes which indicates they may have been acquired during mammalian host adaptation. Several of the unique genes that were potentially acquired show mixed taxonomy across groups indicating convergent evolution by HGT (Extended Data Figure 6 Ldh gene tree, Extended Data Figure 7, nrdD gene). Enolase is thought to be universally present in bacterial genomes but is only found in the oral groups.

Merhej and colleagues^31^ previously reported on the functional differences in COGs between free-living bacteria and many eukaryotic host-dependent bacteria, as well as between free-living and eukaryotic, obligate intracellular bacteria, mutualists and parasites. We performed similar analyses with the Saccharibacteria and the above CPR genomes (Supplemental Table S5) and investigated more closely how organisms with this epibiotic lifestyle compares bacterial mutualists and bacteria considered parasites of eukaryotic hosts. Inter-phylum gene functions show clustering of related groups and functional separation of the G5 G3 and G6 from the G1 group and the G3 oral from rumen (Extended Data Figure 4, Supplemental Table S5). Differences in relative percentages of genes involved with cell wall/membrane/envelope biogenesis, replication recombination and repair, and defense mechanisms are seen in for this unique phylum. Oral G1-G6 groups were low in nucleotide transport and metabolism compared to G1 environmental genomes, mutualists and parasites. Comparisons using SEED families that are more functionally resolved were also compared among the Saccharibacteria groups and select CPR (Extended Data Figure 4). The average GC content also provides clues since this range has also now expanded beyond the average GC content of the G1 genomes (%GC 47 +/- 3%) with G5 the highest at 51%, G3 (40% oral 43% rumen) at a slightly lower range and G6 (32%) representing the lowest of the phylum. It should be noted that the GC content variation however is not entirely consistent with genome reduction process seen in other small, eroded genomes of obligate endosymbionts in restricted host compartments that tend to be AT rich.

The phylogeny indicates that there was a shared ancestral precursor to all the clades including G2-G6 confirming they are related as a phylum. We now know that G3, G5, (and G6) seem exclusively restricted to host associated members (based on SSU data Extended Data Figure1, all have similar reduced metabolic capacity and are small in size. The G-1 group however is the only group to include both mammalian and environmental relatives. From genomic comparisons it is evident that there is some genome reduction occurring from the environment to human oral cavity but synteny is in fact highly maintained in ~70% (first 500Kb the genome) ^1^. The G1 group composed of both environmental and oral representatives is marked by the highest amino acid identities, shared genes (Figure 4, Extended Data Figure 3) suggesting the introduction of G1 from the environment to the host oral cavity was a singular event leading to G1 diversity however we hypothesize that this event was distinct from the human acquisition of G3, and likely G5 group. This represents a sticking point where it is possible that genome reduction occurred independently from the shared environmental ancestor to mammalian hosts in a convergent manner or alternatively a single acquisition event occurred and expansion of the genomic diversity occurred just within the human oral cavity albeit with an already highly reduced genome. Diversity generating retroelements (DGRs), which guide site-specific protein hypervariability are enriched in reduced genomes of some CPR and DPANN archaeal superphylum however the environmental Saccharibacteria genomes investigated are not enriched and contain only 1 DGR per genome ^32^. Although a DGR was found in the oral G1 group TM7x genome, no additional DGRs containing a reverse transcriptase were found in the G1-G6 genomes to help explain their protein diversification. The G3 group is a definitive outlier in this discussion since it clearly shares relatives in mammalian rumen by comparing our G3 rumen genome and 16S rRNA sequence data in current databases from other ruminant mammals including cows, muskox, goat, bighorn sheep (Extended Data Figure 1). This indicates that acquisition of G3 by humans and ruminant mammals occurred within in a shared eukaryotic ancestor or possible lateral transfer between these mammals and humans sometime in the past. Noting that the closest clade to the G3 is the G4 group and members of this G4 are found in sea mammal oral cavities suggest a possible shared non-human ancestor. The distinct genomic similarity in the ruminant and human G3 groups (62% AAI with 56% shared orthologous genes) indicates however a distinct a more recent acquisition in humans than the G1 and G5 groups is the most likely scenario. Mapping of the dental calculus of Neanderthals to oral G3 indicates that less than 1% mapped for this sample which may or may not support if it is indeed not present but it is significantly higher for medieval calculus samples (11-15%) potentially indicating a recent acquisition event during human domestication of ruminant animals although this is just speculation until more data is obtained. For the G5 group, the earlier branching and the overall genomic variation found in the G5 (GC content varies, most unique number of genes of the oral groups) with a lack of close environmental or other mammal representatives indicates that acquisition was also a separate event. At this time there is a complete lack of 16SrRNA for G5 outside of this human group however, if there was a possible unique shared host prior to humans certainly remains an open question.

To further support or refute the possibility of distinct or a single acquisition of the groups, the shared genes in the oral genomes were investigated for evidence of more recent gene transfer by investigating the closest taxonomic hits for each gene and further gene centric phylogenetic investigations. Several unique genes that are shared only among the (HA) groups and are not found in the environmental genomes indicate specific adaptations acquired for optimal growth and survival (Figure 3 panel d). The taxonomic best hits for these select 21 genes unique to the oral cavity (Figure 2) and phylogenetic placement within the domain Bacteria indicate some may have been independently acquired and therefore these lineages converged functionally prior to or within their hosts. All the HA groups have an anaerobic version of the ribonucleoside-triphosphate reductase gene (*nrdD*) and would enable function under low oxygen conditions in the oral cavity and rumen. In addition, these HA groups G3 (oral and rumen) and G5 groups also have *nrdG* gene which enables activation of anaerobic ribonucleoside-triphosphate reductase under anaerobic conditions by generation of an organic free radical, using S-adenosylmethionine and reduced flavodoxin as cosubstrates to produce 5′-deoxy-adenosine. A homolog also exists in the G1 oral group but is more distant (18 to 20% ID) and is annotated as a *pflA*. There is an adenosylcobalamin-dependent ribonucleoside-triphosphate reductase found in RAAC3 environmental genomes and other CPR but is very distant from the anaerobic version (15% ID). The *nrdD* genes between the groups G3 and G5 have different taxonomic hits and are phylogenetically clustered together with other Firmicutes from Clostridia family but apart from G6 which fall within a cluster of predominately Firmicutes bacilli family are also distant (Extended Data Figure 5). By comparison, the TM7x and S2 G1 *nrdD* genes fall within a group of proteobacteria and Elusimicrobia. Another gene unique to the groups (G5, G3 oral and rumen, and G1 TM7x) within the oral cavity and missing completely in environmental G1 genomes was lactate dehydrogenase (ldh). This gene is a signature gene found in many oral bacteria and enables utilization of a common metabolite found in high abundance in plaque. Here, a similar mixture of possible horizontal transfer from disparate groups was found for each of the genes in G5, G3 and G1 (Extended Data Figure 6). Globally, the top taxonomic hits to each of the genes in the genomes (Extended Data Figure 7) varies slightly across the groups and cluster according to their phylogeny for the most part but with interesting divergence of certain groups (Extended Data Figure 7c).

### CRISPR cas system divergence within the Phylum

CRISPR systems were thought to be missing in most CPR ^33^ however we discovered cas genes not previously reported in the oral Sacccharibacteria (Extended Data Figure 8). These include cas1, cas 2 and cas 9. Genome G1 12alb has a ecoli-like Class 1-E Csn2 type gene arrangement (csn2-cas2-cas1-cas9) architecture which is class 2 A(II-A) represented by streptococcus thermophilus CRISPR-associated proteins (cas), as part of the NMENI subtype of CRISPR/Cas loci. The species range so far for this subtype is animal pathogens and commensals only. This protein is present in some but not all NMENI CRISPR/Cas loci. No shared DR or spacers were observed between any of the genome groups.

### Additional shared and unique features of Saccharibacteria groups

Genes maintained in the core pangenome (Supplementary Table S6) include type II secretion biosynthesis machinery that occurs within two regions of the reduced genomes of the G1 and has also been observed in vibrio genomes. We previously reported that TM7x had a large percentage of transmembrane domains (~30%) and a relatively small proportion of coding regions (<3%, 30 CDS) predicted to have signal peptides targeting them for secretory machinery ^1^, this trend also holds true. We found part of the syntenic region of the G1 group includes a novel region that has an array of small hypothetical proteins (4-6) all containing N-terminal signal peptides (Extended Data Figure 9). This unique array of proteins was also found in G3, G5 and G6 genomes as well as other Saccharibacteria and the amoebae symbiont belonging to the lineage TM6 ^34^ (proposed Phylum name Dependentiae). The closest homology of the proteins in this array to any known sequence indicates it may be related to VIRB2 type IV secretion system (Extended Data Figure 9). Further homology search revealed hits to VIRB4 VIRB6 which span the cell membrane in gram positive bacterial envelopes as well as type IV related protein Pgrl and a known DNA-utilizing gene (Gntx) which enables the use of DNA as a substrate in other bacteria. Type IV secretion is involved in transporting a wide range of components, from single proteins to protein-protein and protein-DNA complexes, conjugal transfer of plasmid DNA, and has been shown in the pathogen *Bartonella* to directly inject effectors into host cells ^35^. We hypothesize that duplicated Type II secretion apparatus as well as this type IV-like secretion system in Saccharibacteria and other CPR may be involved in translocation of nutrients from their host to which they are reliant upon. Unique genes in each genome are dominated by hypothetical genes (Supplementary Table S7). However, the functions of the annotated genes will hopefully reveal strategies for cultivation and ideally possible clues of likely hosts although we do not see taxonomic or functional signatures in the TM7x genome that indicate their growth is supported by their already known *Actinomyces odontotlyticus* host.

## Concluding Remarks

In this study we systematically captured the spectrum of known diversity in this phylum by reconstructing completely novel Class level genomes belonging to the highly sought after groups G3, G6 and G5 from humans and also G3 from mammalian rumen. Additional genomes for representatives of G1 were also obtained from modern oral plaque and ancient dental calculus. Overall the genomes were similar to other known Saccharibacteria in that all shared reduced genomes and reduced capacities for many *de novo* biosynthetic pathways further supporting their likely epibiotic lifestyle similar to the cultivated TM7x strain. Global comparative phylogenetic and phylogenomics along with genomic, functional and taxonomic analysis at the gene level highlighted the diversity across this phylum. Within the human oral cavity alone, variation in as much as 70% of the genes from nearest oral clade (AAI 50%) as well as wide GC content variation is evident in these newly captured divergent members (G3, G5 and G6) with no environmental relatives. We found that the human oral G3 shares a close relative in mammalian rumen which is also supported in the 16S rRNA sequences deposited from other ruminant mammals. Independent episodes of transmission of in each of these TM7 groups into humans from environmental and possibly host associated ruminant mammals is likely given the co-evolutionary changes seen within groups and convergent evolution of several key functions during adaptation within hosts. We further provided evidence from *in vivo* collected samples that each of these major groups are ultra-small in size and are found attached to larger cells, yet much work in cultivation is needed to ultimately identify their supporting host organism and if indeed they also share an epibiotic mode of growth and also the host killing parasitic tendencies of strain TM7x.

## Acknowledgements

This research was funded by NIH NIGMS R01GM095373 (J.S.M.), 1R01DE023810 (W.S., X.H., and J.S.M.), 1R01DE020102 (W.S., X.H., and J.S.M.), 1R01DE026186 (W.S., X.H., and J.S.M.).

**Availability of data and material**: The genomes will be made available on NCBI under the Bioproject PRJNA384792 upon acceptance of this manuscript and via https://research.dental.uw.edu/mclean/downloads/.

## Methods

### Genome recovery of novel groups from In vitro growth enrichments and publically available metagenomic datasets

Supplemental Table 1 describes where the assembled genomes were derived either from existing publically available assemblies with reference ID’s (some manually curated, see description below), binned from metagenomics reads (public or from in-house sequencing). MegaBLAST against HOMD 16S rRNA database V14.5 ^1^ was used to find assemblies with hits to TM7 sequences within the 1100 HMP assemblies. Raw reads from in-house sequencing of enrichment samples or public SRA files were assembled after quality-trimming and filtering via BBDuk ^2^ and assembly with St. Petersburg genome assembler (SPAdes) ^3^. We used CONCOCT ^4^ for binning scaffolds. Manual curation of binned genomes was performed by screening for contigs with duplicated genes, GC content and for contigs that had deviating Kmer frequencies validated visually with VizBin in an iterative manner ^5^. Manual curation of publically available draft genomes using Kmer frequency check (IMG tool) was performed before further analyses and contaminating contigs were manually removed from the assembly (See Supplementary Table S1). The resulting assemblies listed in the tree within Figure 1 and Supplementary Table S1. In addition, CheckM was used to assess the quality and estimate the coverage of a set of single marker genes for each genome ^6^. To phylogenetically place the Saccharibacteria, 15 assemblies and 3 publically available CPR genomes (SR1, Kazan, Berkelbacteria from Brown et al. ^7^; Supplemental Table S1) used as outgroups. PhyloSift ^8^ and the associated genome database as well as CheckM was used to extract align and concatenate single copy ribosomal marker genes from assemblies representing genomes/bins. Concatenated alignments were masked to include 90% of the columns (columns with 10% gaps were removed). Concatenated trees were inferred using RAXML (version 8.2.7, GTR CAT rapid bootstrap and search for best scoring tree; bootstrap replicates 100. Parsimony random seed 1 with 800 columns. Likelihood of final tree evaluated and optimized under GAMMA model) ^9^. For phylogenomic placement, both PhyloSift which places the concatenated alignments onto a larger tree and CheckM, which only generates a concatenated alignment using the genome set as input, were congruent in the placement of the Saccharibacteria genomes relative to each other. Individual protein alignments of to investigate phylogenic origins of potentially horizontally acquired genes were also aligned, masked and trees inferred with RAXML with same settings as the concatenated trees. Phylogenetic trees were visualized and annotated with FigTree v1.2.2 ^10^.

### Annotation and metabolic reconstruction

Open reading frames (ORFs) were predicted and annotated for all previously published genomes and genomic scaffolds using Prokka to enable genomics comparisons ^11^. CompareM was employed to perform comparative genomic analyses including the calculation of pairwise amino acid identity (AAI) values between genomes ^12^. The Saccharibacteria phylum pangenome was determined using BLAST-based all vs all comparisons with Prokka-annotated genomes using Roary ^13^ (minimum identity set to 20), which clusters proteins using MCL-edge ^14^. Reconstruction of gene content evolution in Saccharibacteria was performed using COUNT software ^15^ to infer gene gains and losses on the concentrated marker tree in Figure 1 from the matrix of phyletic patterns (containing presence/absence information for each genome in an orthologous group out of 2905 groups). Parameter optimization for a phylogenetic birth-and-death model was employed and Dollo parsimony was used to infer gene gain and loss for these genomes relying on the previously inferred states at ancestral tree nodes. GhostKOALA webserver ^16^ was used to derive taxonomy for each gene in the genomes. Each query gene was assigned a taxonomic category according to the best-hit gene in the Cd-hit cluster supplemented version of their non-redundant dataset. EggNOG-mapper ^17^ was used on protein sequences to determine COG categories. The percentages of hits to each COG category divided by the total COG hits per genome and compared with previously reported data ^18^. ClustVis ^19,20^ was used to make heatmaps and PCA plots of the COG data.

### SSU analysis

For SSU analysis, 16S rRNA genes extracted from genomes that contained a full length sequences for this gene and the HOMD full length 16S rRNA genes from the TM7 phylum Groups 1-6 were aligned with reference sequences with the SINA aligner ^21^ using the SILVA web interface ^22^ with default parameters. Nearest neighbors references (n-20; 89% cutoff) were extracted for each sequence. Unique sequences from this set were extracted in Geneious version 11 (http://www.geneious.com) ^23^. The resulting 400 sequences in the SILVA alignment were masked as above with >10% gaps removed. Trees were inferred by RAxML, as described above (Figure 1 and Extended Data Figure 1). The complete 16S rRNA tree is available in nexus format as Supplemental Data 1.

### Read mapping to new reference genomes

Warinner et al sequenced the dental calculus using whole DNA shotgun methods from adult human skeletons found in a Medieval monastic site ^24^. Reads from 4 samples (7 to 13M reads) were mapped to the reference TM7 genomes. Ancient, dental calculus was also deeply sequenced (>147 million reads) from the well-preserved Neanderthal, El Sidrón 1, which suffered from a dental abscess ^25^ as well as more degraded samples from SPYII. Percentage of genome coverage across these datasets as well as periodontal disease samples,^26^, and HMP datasets (Accession numbers indicated in Supplementary Table S4) were used to assess presence of near relatives in the samples chosen (Figure 2 and Supplementary Table S4).

### Functional comparisons

To compare the encoded functions of Saccharibacteria and select CPR with other known free-living bacteria and those that are host-dependent (eukaryotic host-dependent bacteria and obligate intracellular bacteria) the genes within Clusters of Orthologous Groups (COGs) were determined using EggNOG-mapper ^17^. This comparison is limited to genomes of bacteria that interact with eukaryotic hosts since very limited information is available for genomes of bacteria that interact with bacterial hosts such as TM7x. Merhej and colleagues ^18^ determined the mean number of genes assigned to each COG for bacterial with different lifestyles. They compared the mean number of genes assigned to each COG function between free-living bacteria (125 organisms) and all eukaryotic host-dependent bacteria (125 organisms), then between free-living and eukaryotic and obligate intracellular bacteria (40 organisms), and between mutualists (13 organisms) and parasites (27 organisms). We performed the same analyses with the new CPR genomes to determine how its functional repertoire at the COG level compares to sequenced bacterial mutualists and other bacteria considered parasites of eukaryotic hosts.

### FISH staining of saliva samples and bacteria samples collected from saliva or tongue surfaces

Human oral saliva and tongue-surface bacterial samples were collected fresh. Samples were centrifuged at 1500 rpm for 3 minutes to remove large food particles and other debris. The supernatant was collected and centrifuged at 4600 rpm for 15minutes and the pellets were resuspended and fixed in 4% formaldehyde for 3 hours and permeabilized by 2 mg/mL lysozyme in 20 mM Tris pH7.0 for 9 minutes at 37°C. Fixed cells were washed in 50, 80 and 90% of ethanol, and resuspended in 100-300 µL of hybridization buffer (20 mM Tris•Cl, pH8.0, 0.9 M NaCl, 0.01% SDS, 30% deionized formamide) and incubated at 37°C for 30 minutes. TM7-clade or SR1 specific probes (see table below) were used to stain the cells for 3 hours at 42°C. Cells were then washed three times with 0.1x saline-sodium citrate buffer, 15 minutes each, and mounted on the cover slip with SlowFade Gold antifade reagent (Invitrogen). During the second wash step, Syto9 universal DNA stain (Invitrogen) was added to the samples. Cells were visualized with a LSM 880 inverted confocal microscope equipped with an100x/1.15 oil immersion objective. We repeated each experiment multiple times and acquired multiple FISH images, and only representative images are shown. To insure the clade-specificity of the DNA probes, we performed FISH analysis on TM7x (G1) strain ^27^ using different TM7-clade specific probes, and showed that all the probes, except the G1-specific probe, do not stain TM7x cells (see table below).

We screened for the presence of TM7-clades in 12 individuals using PCR. Two individuals showing positive identification for TM7-G3, G5, G6 and SR1 were studied further. 10 mL of saliva samples were collected from each of the 2 subjects in the morning before brushing their teeth. Saliva samples were mixed 1:1 with cold PBS and kept on ice. Samples were vortexed vigorously for 10 minutes and then centrifuged for 15 minutes at 2600 x g. Pellets were discarded and the supernatant was filtered through 41 µm filter to further remove large particles. Resulting samples were filtered through 0.45 µm filter and centrifuged at 120, 000 x g for 90 minutes at 4°C to concentrate the ultra-small bacteria. The supernatant was discarded and the pellet was resuspended in 300 µL of PBS. The genomic DNA was isolated from these samples and the presence of specific clade of TM7 and SR1 was detected by PCR using clade-specific primers (see table below). The PCR reaction mixture (25 µL) contained 0.5 mM primers, buffer and Taq polymerase. The reactions were incubated at an initial denaturation at 95 °C for 5 min, followed by a 30-cycle amplification consisting of denaturation at 95 °C for 1 min, annealing at different temperature for varying duration depend on different primer sets (see table below), and extension at 72 °C for 2 min.

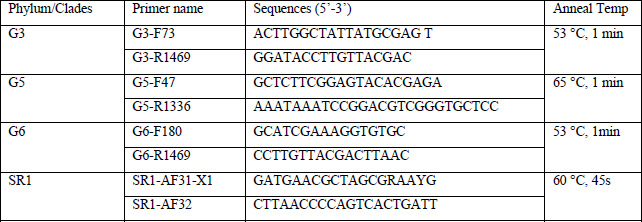

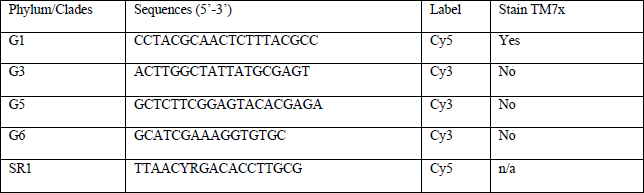

